# Mapping mutational effects along the evolutionary landscape of HIV envelope

**DOI:** 10.1101/235630

**Authors:** Hugh K. Haddox, Adam S. Dingens, Sarah K. Hilton, Julie Overbaugh, Jesse D. Bloom

## Abstract

The immediate evolutionary space accessible to HIV is largely determined by how single amino-acid mutations affect fitness. These mutational effects can shift as the virus evolves. However, the prevalence of such shifts in mutational effects remains unclear. Here we quantify the effects on viral growth of all amino-acid mutations to two HIV envelope (Env) proteins that differ at >100 residues. Most mutations similarly affect both Envs, but the amino-acid preferences of a minority of sites have clearly shifted. These shifted sites usually prefer a specific amino acid in one Env, but tolerate many amino acids in the other. Surprisingly, shifts are only slightly enriched at sites that have substituted between the Envs -- and many occur at residues that do not even contact substitutions. Therefore, long-range epistasis can unpredictably shift Env's mutational tolerance during HIV evolution, although the amino-acid preferences of most sites are conserved between moderately diverged viral strains.

## Introduction

HIV's envelope (Env) protein evolves very rapidly. The major group of HIV-1 that is responsible for the current pandemic originated from a virus that entered the human population ∼100 years ago (***Sharp and Hahn, 2011; Worobey et al., 2008; Faria et al., 2014***). The descendants of this virus have evolved so rapidly that their Envs now have as little as 65% protein identity (***Lynch et al., 2009***). For comparison, protein orthologs shared between humans and mice have only diverged to a median identity of 78% over 90 million years (***Waterston et al., 2002; Hedges et al., 2006***).

Env's rapid evolution has dire consequences for anti-HIV immunity, since it erodes the efficacy of most neutralizing antibodies (***Albert et al., 1990; Wei et al., 2003; Richman et al., 2003; Burton et al., 2005***). Because of this public-health importance, numerous studies have experimentally characterized aspects of the "evolutionary landscape" that Env traverses. The immediate evolutionary space accessible to any given Env is largely defined by the effects on viral fitness of all single amino-acid mutations to Env. Most mutational studies have measured how just a small number of these mutations affect viral growth in cell culture, although it has recently become possible to use deep mutational scanning to measure the effects of many (***Al-Mawsawi et al., 2014; Duenas-Decamp et al., 2016***) or even all (***Haddox et al., 2016***) single amino-acid mutations mutations to an Env variant.

But interpreting these studies in the context of Env evolution requires addressing a fundamental question: How informative are mutational studies of a single protein variant about constraints on long-term evolution? During protein evolution, substitutions at one site can change the effect of mutations at other sites (***Natarajan et al., 2013; Gong et al., 2013; Harms and Thornton, 2014; Podgornaia and Laub, 2015; Starr and Thornton, 2016; Klink and Bazykin, 2017***). We will follow the nomenclature of ***Pollock et al. (2012)*** to refer to these changes in mutational effects as *shifts* in a site's amino-acid preferences. Such shifts can accumulate as substitutions become entrenched via epistatic interactions with subsequent changes (***Starr et al., 2017; Pollock et al., 2012; Shah et al., 2015; Bazykin, 2015***) - although the magnitude of these shifts is usually limited (***Doud et al., 2015; Chan et al., 2017; Ashenberg et al., 2013; Risso et al., 2014***).

Given that the Envs of circulating HIV strains represent a vast collection of homologs that often differ at >100 residues, shifts in amino-acid preferences could make the outcome of any study highly dependent on the Env used. The extent to which this is actually the case is unclear, since the few protein-wide studies of such shifts have examined proteins that are structurally far simpler than Env, which forms a large heavily glycosylated heterotrimeric complex that transitions through multiple conformational states (***Munro et al., 2014; Ozorowski et al., 2017***).

Here we use an improved version of a previously described deep mutational scanning strategy (***Haddox et al., 2016***) to measure the effects on viral growth of all single amino-acid mutations to two transmitted-founder virus Envs that differ by >100 mutations. We compare these complete maps of mutational effects to identify sites that have shifted in their amino-acid preferences between the Envs. Most sites show no detectable shifts, but 30 sites have clearly shifted preferences. These shifted sites usually prefer a specific amino acid in one Env but have shifted to tolerate many amino acids in the other Env. The shifted sites cluster in structure, but are often distant from any amino-acid substitutions that distinguish the two Envs, demonstrating the action of long-range epistasis. By aggregating our measurements for both Envs, we identify sites that evolve faster or slower in nature than expected given the functional constraints measured in the lab, probably due to pressure for immune evasion. Overall, our work provides complete across-strain maps of mutational effects that inform analyses of Env's evolution and function.

## Results

### Two Envs from clade A transmitted-founder viruses

The viruses most relevant to HIV's long-term evolution are those which are transmitted from human-to-human. However, the only prior work that has measured how all Env amino-acid mutations affect HIV growth is a study by some of us (***Haddox et al., 2016***) that used a late-stage lab-passaged CXCR4-tropic virus (LAI; ***Peden et al., 1991***). The properties of Env can vary substantially between such late-stage viruses and the transmitted-founder viruses relevant to HIV's long-term evolution (***Sagar et al., 2006; Wilen et al., 2011; Parrish et al., 2013; Ronen et al., 2015***).

We therefore selected Envs from two transmitted-founder viruses, BG505.W6M.C2.T332N and BF520.W14M.C2 (hereafter referred to as BG505 and BF520), that were isolated from HIV-infected infants shortly after mother-to-child transmission (***Nduati et al., 2000; Wu et al., 2006; Goo et al., 2014***). The BG505 Env has been extensively studied from a structural standpoint (***Julien et al., 2013; Lyumkis et al., 2013; Pancera et al., 2014; Huang et al., 2014; Sanders et al., 2015; Stewart-Jones et al., 2016; Gristick et al., 2016***), and variants of this Env are being tested as a vaccine immunogens (***Sanders et al., 2013,2015; de Taeye et al., 2015***). We used the T332N variant of BG505 Env because it has a common glycosylation site that is targeted by many anti-HIV antibodies (***Sanders et al., 2013***). The BF520 Env was isolated from an infant who developed an early broad anti-HIV antibody response (***Goo et al., 2014; Simonich et al., 2016***). We have previously created comprehensive codon-mutant libraries of the BF520 Env and used them to map HIV antibody escape (***Dingens et al., 2017***), but these BF520 libraries have not been characterized with respect to how mutations affect viral growth.

Both BG505 and BF520 are from clade A of the major (M) group of HIV-1. Figure 1 shows the phylogenetic relationship among these two Envs and other clade A sequences. BG505 and BF520 are identical at 721 of the 836 pairwise-alignable protein sites (86.2% identity). In our experiments, we analyze only the ectodomain and transmembrane domain of Env (we exclude the signal peptide and cytoplasmic tail). In these regions of Env, BG505 and BF520 are identical at 549 of the 616 sites (89.1% identity) that are alignable across clade A Envs (Figure 1-source data 1, Figure 1-source data 2). The divergence between BG505 and BF520 therefore offers ample opportunity to investigate mutational shifts during Env evolution.

**Figure 1.**
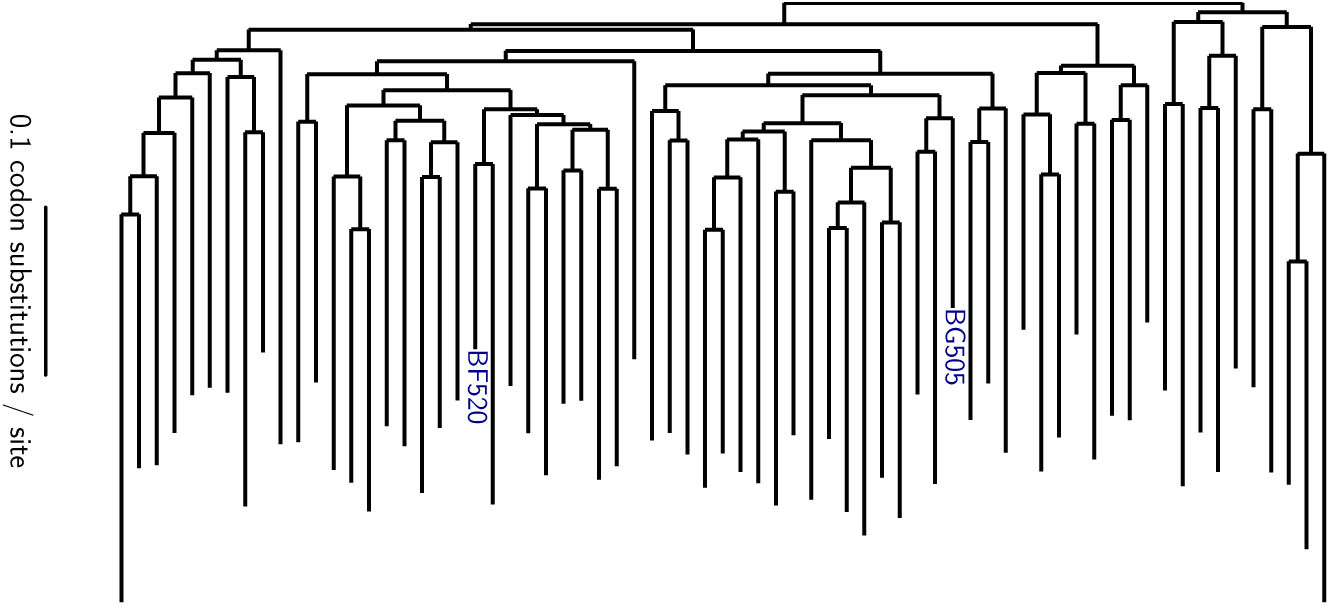
Phylogenetic tree showing the relationship of BG505 and BF520 to other clade A Envs. The tree shows the 69 Envs in the alignment in Figure 1-source data 1, which is a subsample of clade A sequences from the group M alignment in the Los Alamos HIV sequence database (http://www.hiv.lanl.gov). Sites not mutagenized in our experiments (the signal peptide and cytoplasmic tail) or that are poorly alignable were masked as indicated in Figure 1-source data 2, leaving 616 alignable sites. The pairwise identity of BG505 and BF520 to other sequences at alignable sites is in Figure 1-Figure supplement 1. The tree topology was inferred using RAxML (***Stamatakis, 2014***) under the GTRCAT model of nucleotide substitution, and branch lengths were optimized under the M0 Goldman-Yang model (***Yang et al., 2000***) using phydms (***Hilton et al., 2017***). **Figure 1-Figure supplement 1**. Pairwise identity of all Env sequences to BG505 and BF520. **Figure 1-source data 1**. The alignment of clade A ***env*** coding sequences is in `cladeA_alignment.fasta`. **Figure 1-source data 2**. The 240 Env sites masked in all phylogenetic analyses because they were not mutagenized in our experiments or are poorly alignable are listed in `alignment_mask.csv`.

### Deep mutational scanning of each Env

We have previously described a deep mutational scanning strategy for measuring how all amino-acid mutations to Env affect HIV growth in cell culture, and applied this strategy to the late-stage lab-adapted LAI strain (***Haddox et al., 2016***). Here we made several modifications to this earlier strategy to apply it to transmitted-founder Envs and to reduce the experimental noise. This last consideration is especially important when comparing Envs, since it is only possible to reliably detect differences that exceed the magnitude of the experimental noise. Our modified deep mutational scanning strategy is in Figure 2A. This approach had the following substantive changes: we used SupT1 cells expressing CCR5 to support growth of viruses with transmitted-founder Envs, we used more virions for the first passage (≥ 3 × 10^6^ versus 5 × 10^5^ infectious units per library) to avoid bottlenecking library diversity, and rather than performing a full second passage we just did a short high-MOI infection to enable recovery of ***env*** genes from infectious virions without bottlenecking (Figure 2A). We performed this deep mutational scanning in full biological triplicate for both BG505 and BF520 (Figure 2B). Our libraries encompassed all codon mutations to all sites in Env except for the signal peptide and cytoplasmic tail.

**Figure 2.**
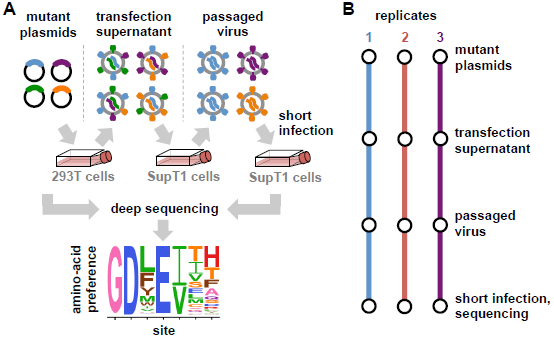
Deep mutational scanning workflow. **(A)** We made libraries of proviral HIV plasmids with random codon-level mutations in the ***env*** gene. The number of mutations per gene approximately followed a Poisson distribution with a mean between 1 and 1.5 (Figure 2-Figure supplement 1). We transfected the plasmids into 293T cells to generate mutant viruses, which lack a genotype-phenotype link since cells are multiply transfected. To establish a genotype-phenotype link and select for Env variants that support HIV growth, we passaged the libraries in SupT1.CCR5 cells for four days at a low multiplicity of infection (MOI) of 0.01. To isolate the ***env*** genes from only viruses that encoded a functional Env protein, we infected the passaged libraries into SupT1.CCR5 cells at high MOI and harvested reverse-transcribed non-integrated viral DNA after 12 hours. We then deep sequenced the ***env*** genes from these final samples as well as the initial plasmid library, using molecular barcoding to reduce sequencing errors. We also deep sequenced identically handled wildtype controls to estimate error rates. Using these sequencing data, we estimated the preference for each of the 20 amino acids at each site in Env. These data are represented in logo plots, with the height of each letter proportional to that site's preference for that amino acid. **(B)** We conducted this experiment in full biological triplicate for both BG505 and BF520, beginning each replicate with independent creation of the plasmid mutant library. These replicates therefore account for ***all*** sources of noise and error in the experiments. **Figure 2-Figure supplement 1.** Sanger sequencing of selected clones from the mutant plasmid libraries.

The deep mutational scanning effectively selected for functional Envs as evidenced by strong purifying selection against stop codons. Figure 3A shows the average frequency of mutations across Env in the plasmid mutant libraries, the mutant viruses, and wildtype controls as determined from the deep sequencing. The mutant viruses show clear selection against stop codons and many nonsynonymous mutations (Figure 3A). This selection is more apparent if we correct for the background error rates estimated from the wildtype controls (Figure 3-Figure-supplement 1). The error-corrected frequencies of stop codons drop to 3%-16% of their original values (Figure 3-Figure-supplement 1), with the residual stop codons probably due to some non-functional virions surviving due to complementation by other co-infecting virions. The error-corrected frequencies of nonsynonymous mutations also drop substantially (43%-49% of their original values), whereas the frequencies of synonymous mutations drop only slightly (85%—95% of their original values). These trends are consistent with the fact that nonsynonymous mutations are often deleterious, whereas synonymous mutations often (although certainly not always, see ***Zanini and Neher, 2013***) have only mild effects on viral growth. Figure 3A only summarizes one aspect of the deep mutational scanning data, but Supplementary files 1 and 2 contain detailed plots showing all aspects of the data (read depth, per-site mutation rate, etc) as generated by the dms_tools2 software (***Bloom, 2015***, https://jbloomlab.github.io/dms_tools2/).

**Figure 3.**
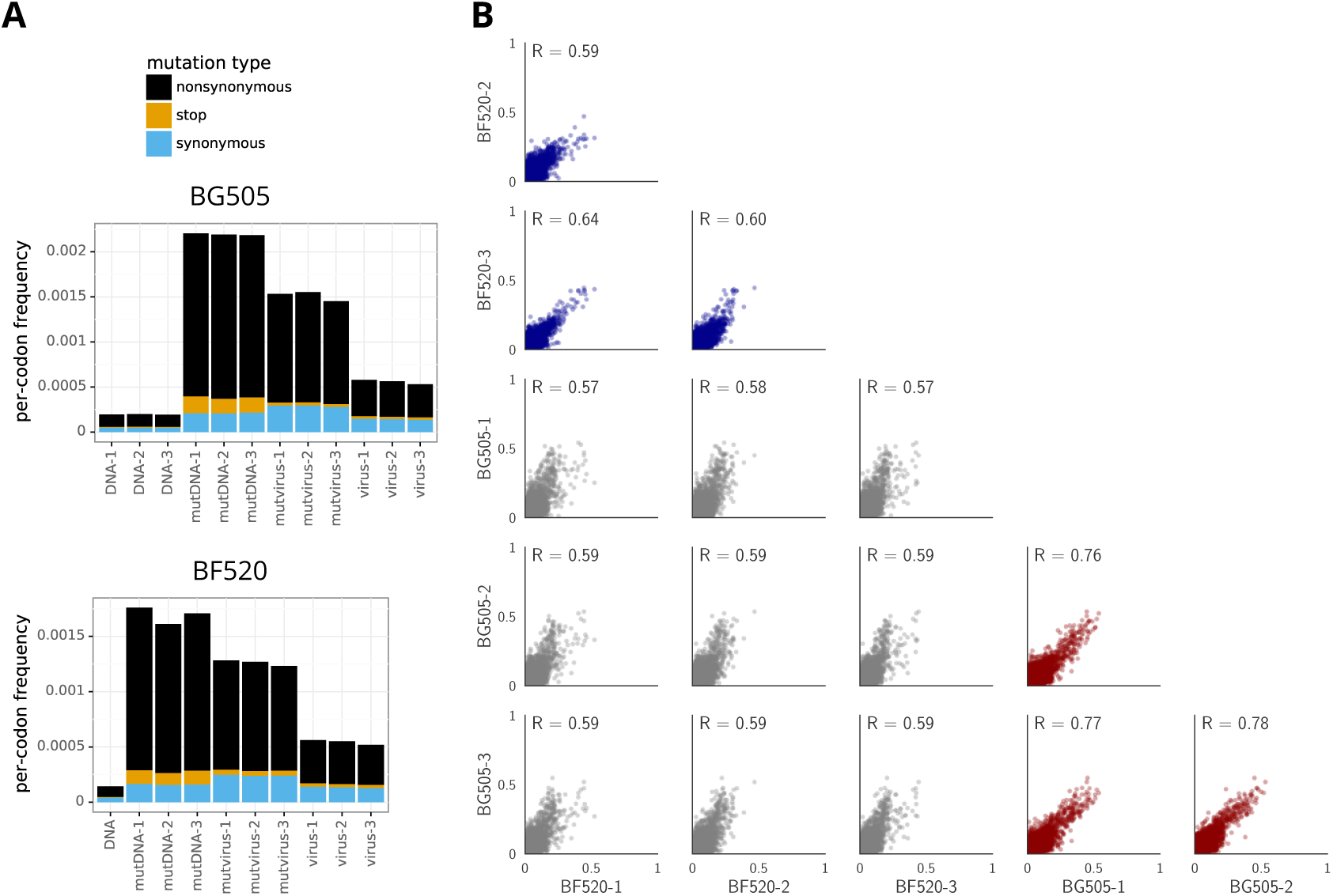
The deep mutational scanning selects for functional Envs and yields measurements that are well correlated among replicates. **(A)** The average per-codon mutation frequency when sequencing plasmids encoding wildtype Env (*DNA*), plasmid mutant libraries (*mutDNA*), mutant viruses after the final infection (*mutvirus*), and virus generated from wildtype plasmids (*virus*). Mutations are categorized as nonsynonymous, synonymous, or stop codon. The *DNA* samples show that sequencing errors are rare, and the *virus* samples show that viral-replication errors are well below the frequency of mutations in the *mutDNA* samples. Comparing the *mutvirus* to *mutDNA* shows clear purifying selection against stop codons and some nonsynonymous mutations, particularly after subtracting the background error rates given by the *virus* and *DNA* samples (Figure 3-Figure supplement 1). More extensive plots from the analysis of the deep sequencing data are in Supplementary files 1 and 2. **(B)** Correlations between replicates in the measured preferences of each site in Env for all 20 amino acids. Blue indicates replicate measurements on BF520' red indicates replicate measurements on BG505, and gray indicates across-Env measurements of BF520 versus BG505. *R* is the Pearson correlation coefficient. The numerical values for the preferences are in Figure 3-source data 1. **Figure 3-Figure supplement 1**. Numerical frequencies of mutations. **Figure 3-source data 1**. Preferences for each replicate and averages are in `all_prefs_unscaled.zip`.

We used the deep mutational scanning data to estimate the preference of each site in Env for each amino acid via the analysis method described in ***Bloom (2015)***. As graphically illustrated in Figure 2A, the preferences for each site are normalized to sum to one. Our libraries were mutagenized at 670 sites in BG505 and 662 sites in BF520, so 670×20 = 13.400 and 662×20 = 13.240 preferences were estimated for each Env, respectively. The correlations between the preferences from different experimental replicates are in Figure 3B. These replicate-to-replicate correlations are substantially higher than those for the deep mutational scanning of LAI Env by ***Haddox et al. (2016),*** which had replicate-to-replicate Pearson correlations of only ***R*** = 0.45 to 0.50.

While the replicates are well correlated across all replicates for both BG505 and BF520, the replicates for BG505 are more correlated with each other than with replicates for BF520, and vice versa (Figure 3B, compare red and blue versus gray plots). This fact hints that there are some shifts in amino-acid preferences between the two Envs - something that is investigated with more statistical rigor later in this paper. Note also that there is a trend for highly preferred amino acids to be more strongly preferred in BG505 than BF520 (most high-preference points in the gray plots in Figure 3B fall above the diagonal); however, this trend does not necessarily reflect differences between the Envs. Rather, there were modest differences in the stringency of selection between our deep mutational scans of BG505 and BF520 (Figure 3-Figure-supplement 1 shows that purifying selection better purged stop codons in BG505). In the next section, we correct for these experimental differences by calibrating each dataset to match the stringency of selection in nature.

### Amino-acid preferences of the Envs and their relationship to HIV evolution

The most immediate question is how authentically the experimental measurements describe the actual selection on Env function in nature. Direct comparisons between experimentally measured amino-acid preferences and amino-acid frequencies in natural sequences are confounded by the fact that the natural sequences are evolutionarily related. This problem can be overcome by making the comparison in a phylogenetic context. Specifically, we used our deep mutational scanning data to construct experimentally informed codon models (ExpCM's) for Env evolution (***Hilton et al., 2017***). Importantly, since we expect many sites in Env to be under diversifying selection from immunity, we extended the ExpCM's to draw the relative dN/dS parameter (ω) from a gamma distribution as is commonly done for codon-substitution models (***Yang et al., 2000***).

Table 1 shows that phylogenetic models informed by the deep mutational scanning of either BG505 or BF520 describe the natural evolution of Env vastly better than a standard codon substitution model. In addition to simply comparing model fits, we can interpret the model parameters. ExpCM's have a stringency parameter that relates selection in the experiments to that in nature. A stringency parameter >1 indicates that natural selection prefers the same amino acids as the experiments, but with greater stringency (***Hilton et al., 2017***). Both ExpCM's have stringency parameters >1 (Table 1) - a finding that makes sense, since the stop-codon analysis in the previous section suggests that the experimental selections are more lax than natural selection on HIV. We can also interpret the *ω* parameter. For standard codon substitution models, *ω* is the rate of fixation of nonsynonymous mutations relative to synonymous ones - and the gene-wide average *ω* is almost always <1, since purifying selection purges many functionally deleterious amino-acid mutations even for adaptively evolving proteins (***Murrell et al., 2015***). Indeed, Table 1 shows that Env's gene-wide average *ω* is <1 for a standard model. But for ExpCM's, *ω* is the relative rate of nonsynonymous to synonymous substitutions *after* accounting for functional constraints measured in the deep mutational scanning (***Bloom, 2017***). Table 1 shows that the ExpCM's have a gene-wide average *ω* that is >1, indicating that external selection (e.g., from immunity) drives Env to fix amino-acid mutations faster than expected under a null model that only accounts for functional constraints.

**Table 1.** Evolutionary models informed by the deep mutational scanning describe HIV's evolution in nature much better than a standard substitution model. Shown are the results of maximum likelihood fitting of substitution models to the clade A phylogeny in Figure 1. Experimentally informed codon models (ExpCM, ***Hilton et al., 2017***) utilizing the across-replicate average of the deep mutational scanning describe Env's natural evolution far better than a standard codon substitution model (the M5 model of ***Yang et al., 2000***) as judged by comparing the Akaike information criteria (AAIC, ***Posada and Buckley, 2004***). Both ExpCM models have a stringency parameter >1. All models draw *ω* from a gamma distribution, and the table shows the mean (ω̅) and shape parameters (*ω_α_* and *ω_β_*) of this distribution. The last two columns show the number of sites evolving faster (*ω_r_* > 1) or slower (*ω_r_* < 1) than expected at a false discovery rate of 0.05, as determined using the approach in ***Bloom (2017)***. Analyses were performed using phydms (http://jbloomlab.github.io/phydms/). Table 1-source-data 1 shows the results for additional substitution models. **Table 1-source data 1.** Results for phylogenetic models where *ω* is not drawn from a gamma-distribution or where the preferences are averaged across sites to eliminate the site specificity are in `modelcomparison.md`.

| Model | $\Delta AIC$ | LogLikelihood | nParams | stringency | $\bar{\omega}$ | $\omega_\alpha$ | $\omega_\beta$ | nsites $\omega_r > 1$ | nsites $\omega_r < 1$ |
| --- | --- | --- | --- | --- | --- | --- | --- | --- | --- |
| ExpCM BF520 | 0.0 | -35218.8 | 7 | 2.8 | 1.4 | 1.0 | 0.7 | 66 | 35 |
| ExpCM BG505 | 269.0 | -35353.3 | 7 | 2.1 | 1.3 | 0.9 | 0.7 | 65 | 53 |
| Goldman-Yang M5 | 3552.2 | -36989.9 | 12 | nan | 0.8 | 0.6 | 0.7 | 14 | 211 |

A more qualitative way to assess if the deep mutational scanning authentically describes selection on Env function is to visually compare the measurements with existing knowledge. Figures 4 and 5 show the across-replicate average of the amino-acid preferences for each Env after re-scaling by the stringency parameters in Table 1. (Note that throughout the rest of this paper, we use preferences re-scaled by these stringency parameters.) At sites of known functional importance, these preferences are usually consistent with prior knowledge. For instance, residues T257, D368, E370, W427, and D457 are important for Env binding to CD4 (***Olshevsky et al., 1990***), and all these amino acids are highly preferred in our deep mutational scanning (Figures 4 and 5). Likewise, Env has 10 disulfide bonds (linking sites 54-74,119-205,126-196,131-157, 218-247, 228-239, 296331, 378-445, 385-418, and 598-604), most of which are important for function (***van Anken et al., 2008***) - and the cysteines at these sites are highly preferred in our deep mutational scanning. The deep mutational scanning is also consistent with prior knowledge about sites that are tolerant of mutations. For instance, Env has five variable loops that mostly evolve under weak constraint in nature (***Starcich et al., 1986; Zolla-Pazner and Cardozo, 2010***) - and most sites in these loops are mutationally tolerant in our deep mutational scanning (see sites indicated by gray overlay bars in Figures 4 and 5, such as 132 to 195). It is beyond the scope of this paper to catalog associations between our measurements and all other prior mutational studies of Env, but the concordance of our findings with the above mutational studies, and the fact that our data improve phylogenetic models of Env's natural evolution, suggest that our measurements authentically describe functional selection on Env.

**Figure 4.**
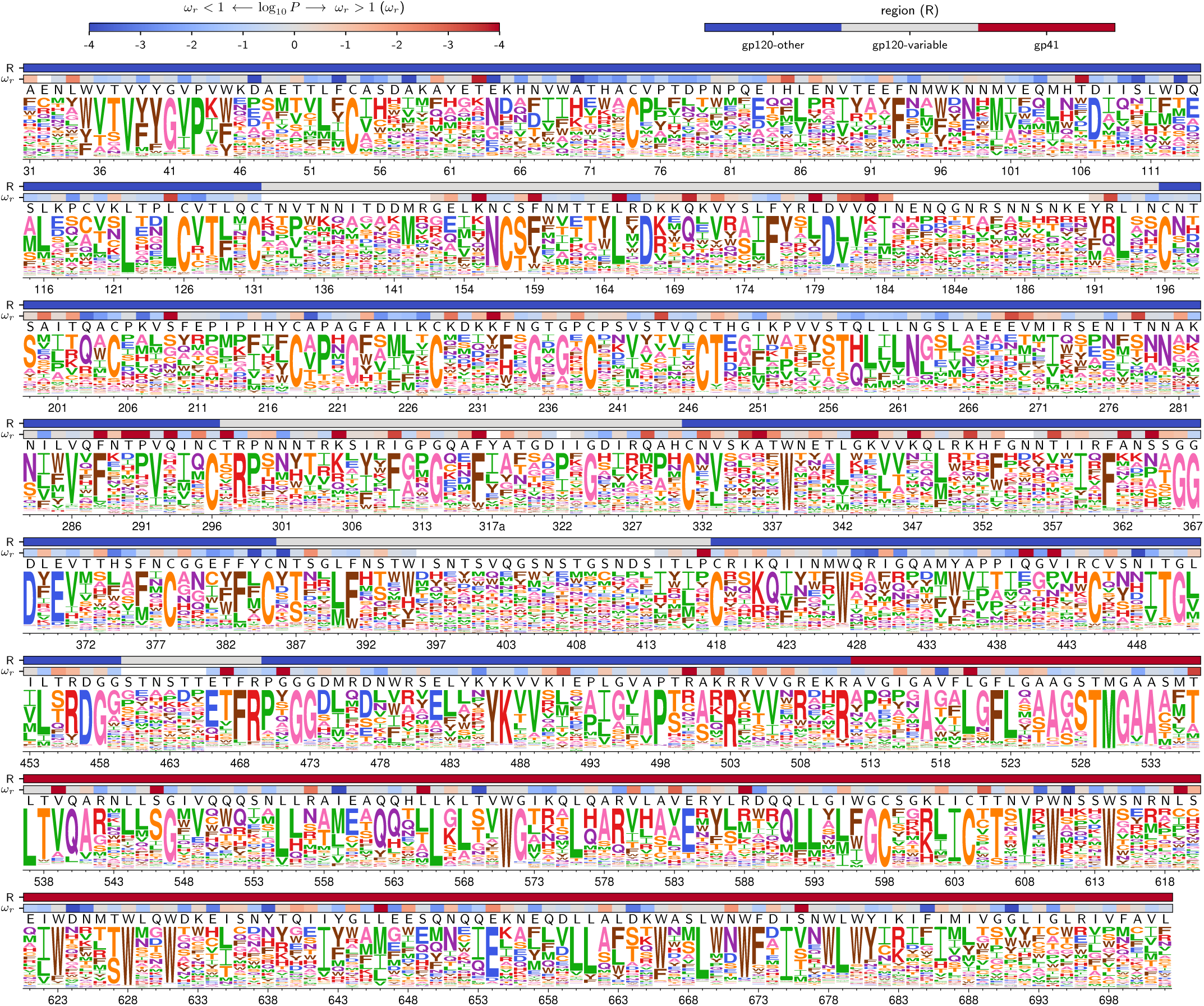
Amino-acid preferences for the BG505 Env. At each site, the height of the letter is proportional for that site's preference for that amino acid. The top color bar indicates the region of Env(gp120 variable loop, gp120 not variable loop, or gp41). The lower color bar indicates the evidence that the site evolves faster (*ω_r_ >* 1) or slower (*ω_r_* < 1) than expected given the experiments (see ***Bloom, 2017***). The letters above the logos indicate the wildtype amino acid in BG505. Sites are numbered using the HXB2 scheme (***Korber et al., 1998***)). This logo plot shows the site-specific amino-acid preferences for BG505 after averaging the replicates and re-scaling by the stringency parameter in Table 1. The figure was generated using dms_tools2 (***Bloom, 2015***, https://jbloomlab.github.io/dms_tools2/)' which in turn utilizes weblogo (***Crooks et al., 2004***). **Figure 4-source data 1**. The numerical values of the amino-acid preferences plotted in this figure are in `rescaled_BG505_prefs.csv`. **Figure 4-source data 2**. The sequence of BG505 Env and mapping from sequential (*original* column) to HXB2 numbering (*new* column) is in `BG505_to_HXB2.csv`. **Figure 4-source data 3**. The *ω_r_* values and associated *P*-values for BG505 in HXB2 numbering are in `BG505_omegabysite.tsv`.

**Figure 5.**
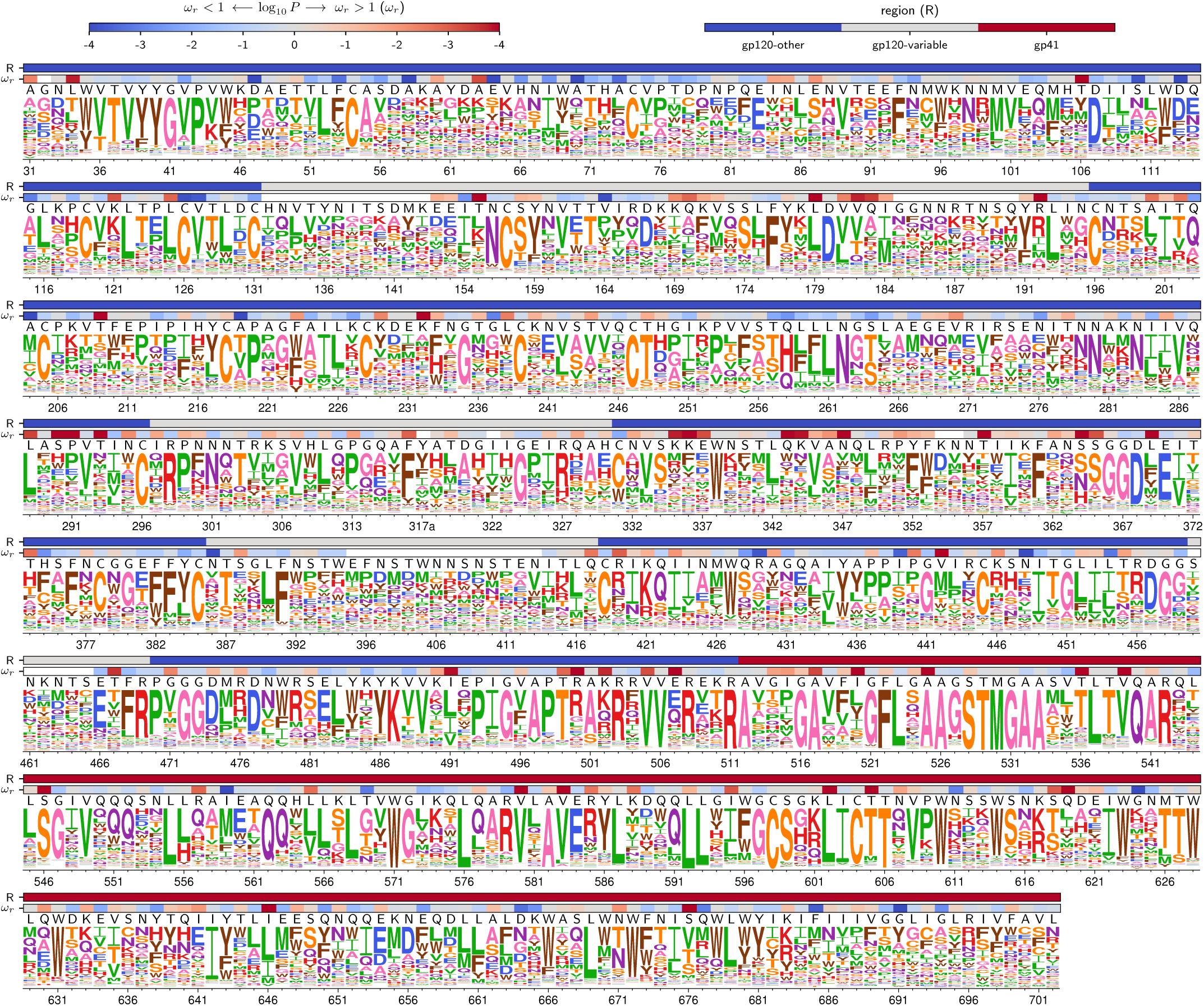
Amino-acid preferences for the BF520 Env. This figure is the same as Figure 4 except that it shows the data for BF520 instead of BG505. **Figure 5-source data 1**. The numerical values of the amino-acid preferences plotted in this figure are in rescaled_BF520_prefs.csv. **Figure 5-source data 2**. The sequence of BF520 Env and mapping from sequential (*original* column) to HXB2 numbering (*new* column) is in `BF520_to_HXB2.csv`. **Figure 5-source data 3**. The *ω_r_* values and associated *P*-values for BF520 in HXB2 numbering are in `BG505_omegabysite.tsv`.

### Shifts in amino-acid preferences between BG505 and BF520

The most fundamental question that we seek to address is how similar the amino-acid preferences are between the two Envs. We have already noted that Figure 3B shows that the preferences are more correlated for replicate measurements on the same Env than for replicate measurements on different Envs. However, simply comparing correlation coefficients does not identify specific sites where mutational effects have shifted, nor does it quantify the magnitude of any shifts.

We therefore used a more rigorous approach to identify sites where the amino-acid preferences differ between BG505 and BF520 by an amount that exceeds the noise in our experiments. We first re-scaled the preferences from each experimental replicate by the stringency parameter for that Env from Table 1 to calibrate all measurements to the stringency of natural selection. We then identified the 659 sites in the mutagenized regions of Env that are pairwise alignable between BG505 and BF520 (Figure 6-source data 1). For each site, we calculated the shift in amino-acid preferences between Envs using an approach similar to that of ***Doud et al. (2015)*** as illustrated in Figure 6A. This approach calculates the magnitude of the shift after correcting for experimental noise by comparing the differences in preferences between replicates for BG505 and BF520 to the differences between replicates for the same Env. Figure 6A shows this calculation for a site that has not shifted (site 598, which strongly prefers cysteine in both Envs), the most shifted site (512, which shifts from being mutationally tolerant in BG505 to strongly preferring alanine in BF520), and two other sites with more intermediate behaviors.

**Figure 6.**
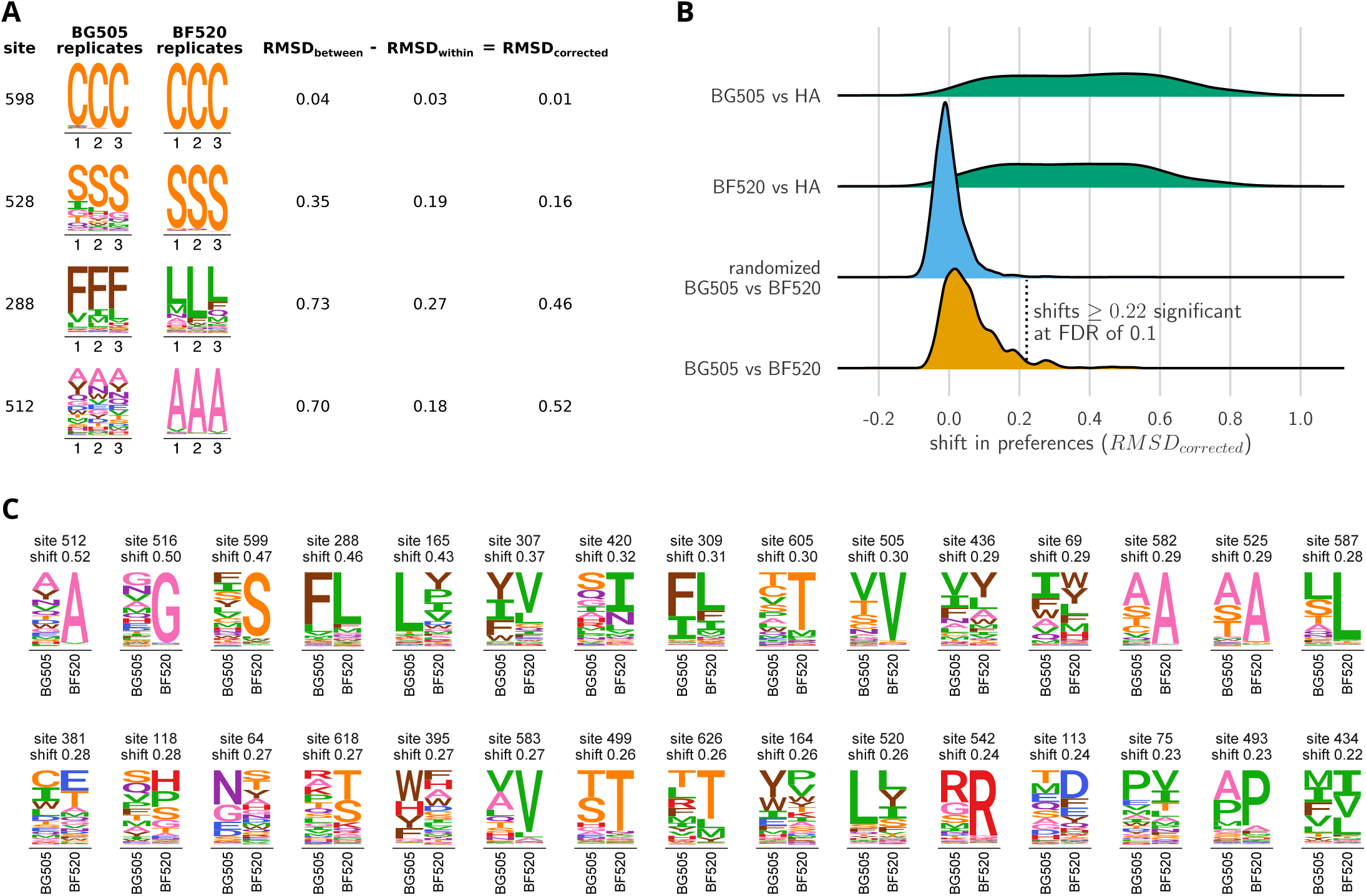
Env sites with shifted amino-acid preferences between BG505 and BF520. (A) Calculation of the corrected distance between the amino-acid preferences of BG505 and BF520 at four example sites. We have triplicate measurements for each Env. We calculate the distance between each pair of replicate measurements, and group these into comparisons *between* the two Envs and *within* replicates for the same Env. We compute the root-mean-square distance (RMSD) for both sets of comparisons, which we denote as RMSD_between_ and RMSD_witllin_. The latter quantity is a measure of experimental noise. The noise-corrected distance between Envs at a site, RMSD_corrected_, is simply the distance between the two Envs minus this noise. **(B)** The bottom distribution (orange) shows the corrected distances between BG505 and BF520 at all alignable sites. The next distribution (blue) is a null generated by computing the corrected distances on all randomizations of the replicates among Envs. The top two distributions (green) compare Env to the non-homologous influenza hemagglutinin (HA) protein (***Doud and Bloom, 2016***) simply putting sites into correspondence based on sequence number. We compute the ***P***-value that a site has shifted between BG505 and BF520 as the fraction of the null distribution that exceeds that shift, and identify significant shifts at a false discovery rate (FDR) of 0.1 using the method of ***Benjamini and Hochberg (1995)***. Using this approach, 30 of the 659 sites have significant shifts (corrected distance ?0.22). **(C)** All sites that have significantly shifted their amino-acid preferences at an FDR of 0.01. For each site, the logo stacks show the across-replicate average preferences for BG505 and BF520. The sites are sorted by the magnitude of the shift. Figure 6-source data 1. The corrected distances between BG505 and BF520 at each site are in `BG505_to_BF520_prefs_dist.csv`.

The overall distribution of shifts between BG505 and BF520 is shown in Figure 6B. Most sites have relatively small shifts (close to zero), although there is a long tail of sites with large shifts. This tail reaches its upper value with site 512, which has a shift of 0.52 out of a maximum possible of 1.0. How should we interpret this distribution - have mutational effects shifted a lot, or not very much? We can establish an upper-bound for how much sites might shift by comparing Env to a non-homologous protein. Figure 6B shows the distribution of shifts when comparing Env to influenza's hemagglutinin protein, which has previously had its amino-acid preferences measured by deep mutational scanning (***Doud and Bloom, 2016***). Most sites have large shifts between Env and hemagglutinin, with the typical shift being ∼0.4 and some approaching the maximum value of 1.0. We can also establish a lower-bound by creating a null distribution for the expected shifts if all differences are simply due to experimental noise. This null distribution is created by randomizing the experimental replicates among Envs. Figure 6B shows that the null distribution is more peaked at zero than the real distribution, and does not have the same prominent tail of sites with large shifts. The answer to the question of how much mutational effects have shifted is therefore nuanced: they have substantially shifted at some sites, but remain vastly more similar between the two Envs than between two unrelated proteins.

We can use the null distribution to identify sites where the shifts between BG505 and BF520 are significantly larger than the noise in our experiments (Figure 6B). There are 30 such sites at a false discovery rate of 0.1. Figure 6C shows the amino-acid preferences of these significantly shifted sites for each Env. For the majority of shifted sites, one Env prefers a specific amino acid whereas the other Env tolerates many amino acids; for instance, see sites 512, 516, 599,165, 605 and 505 in Figure 6C. Such broadening and narrowing of a site's mutational tolerance is frequently linked to changes in protein stability, with a more stable protein typically being more mutationally tolerant (***Wang et al., 2002; Bloom et al., 2006; Gong et al., 2013; Kumar et al., 2017***). Work with engineered Env protein in the form of "SOSIP" trimer (***Binley et al., 2000; Sanders et al., 2002***) has shown that BG505 SOSIP is more thermostable than BF520 SOSIP (***Verkerke et al., 2016***). Consistent with this fact, sites with altered mutational tolerance are often (although not always, see sites 165 and 520 in Figure 6C) more mutationally tolerant in BG505.

However, not all of the significantly shifted sites show a simple pattern of broadening or narrowing of mutational tolerance. For instance, site 288 does not alter its mutational tolerance but rather flips its rather narrow amino-acid preference from phenylalanine in BG505 to leucine in BF520 (Figure 6C). Thus, there is variation in both the extent and types of shifts observed.

### Structural and evolutionary properties of shifted sites

What distinguishes the sites that have undergone significant shifts? Figure 7A shows the locations of the shifted sites on the crystal structure of Env. There is no visually obvious tendency for shifted sites to preferentially be on Env's surface or in its core, and a statistical analysis (Figure 7B) finds no association between a site's relative solvent accessibility and whether its amino-acid preferences have shifted. However, Figure 7A does suggest that the sites of significant shifts tend to cluster in Env's structure, and a statistical analysis confirms that this is the case (Figure 7C). Therefore, the factors that drive shifts in Env's mutational tolerance often affect physically interacting clusters of residues in a coordinated fashion.

**Figure 7.**
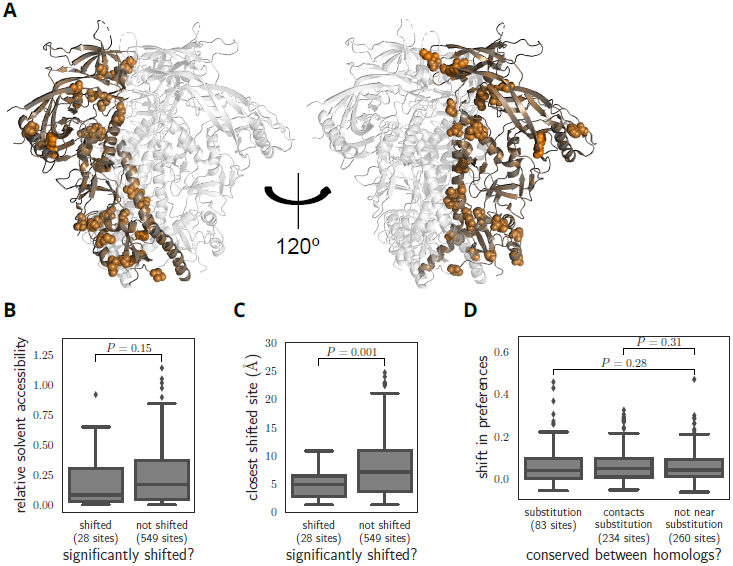
Characteristics of significantly shifted sites. **(A)** One monomer of the Env trimer (PDB 5FYL; ***Stewart-Jones et at., 2016***) is colored from gray to orange according to the magnitude of the mutational shift at each site (orange indicates large shift). Sites that are significantly shifted according to Figure 6B are in spheres, and all other sites are in cartoon representation. **(B)** There is no significant difference in the relative solvent accessibility of sites that have and have not undergone significant shifts. The absolute solvent accessibility of each site was calculated using DSSP (***Kabsch and Sander, 1983***) and normalized to a relative solvent accessibility using the absolute accessibilities from ***Tien et al. (2013)***. **(C)** Sites of significant shifts are clustered in Env's structure. Shown is the distance of each significantly shifted and not-shifted site to the closest other shifted site. **(D)** Large mutational shifts are *not* strongly enriched at sites that have substituted between BG505 and BF520, or at sites that contact sites that have substituted. The plot shows the magnitudes of the shifts among the 83 crystallographically resolved sites that have substituted between BG505 and BF520, the 234 non-substituted sites that physically contact a substitution in the Env structure (any non-hydrogen atom within 3.5Â), and all other sites. Figure 7-Figure supplement 1 shows that there is a modest borderline-significant tendency of significantly shifted sites to have substituted. All panels only show the 577 sites that are resolved in the crystal structure. Of the 30 significantly shifted sites in Figure 6B, only 28 are crystallographically resolved. Structural distances and solvent accessibilities were calculated using all monomers in the trimer. ***P*** values were calculated using the Mann-Whitney ***U*** test. **Figure 7-Figure supplement 1.** Statistical testing of whether significantly shifted sites are more likely to have substituted between BG505 and BF520.

An obvious hypothesis is that strongly shifted sites have substituted between BG505 and BF520, or physically contact such substitutions. According to this hypothesis, substitutions would alter the local physicochemical environment of the substituted site and its neighbors, thereby shifting the amino-acid preferences of sites in the physical cluster. But surprisingly, the typical magnitude of shifts is not significantly larger at sites that have substituted, or at sites that contact sites that have experienced substitutions (Figure 7C). There is a borderline trend for the significantly shifted sites to be more likely to have substituted between BG505 and BF520 (Figure 7-Figure supplement 1), but most shifted sites have not substituted (only 8 of the 30 shifted sites differ in amino-acid identity between the two Envs). The lack of strong enrichment in shifts at substituted sites contrasts with previous protein-wide experimental (***Doud et al., 2015***) and simulation-based (***Pollock et al., 2012; Shah et al., 2015***) studies of shifting amino-acid preferences, which found that shifts were dramatically more pronounced at sites of substitutions. The difference may arise because these earlier studies examined proteins that are fairly conformationally static (absolutely so in the case of the simulations). On the other hand, Env is extremely complex and conformationally dynamic (***Munro et al., 2014; Ozorowski et al., 2017***), which may increase the opportunities for long-range epistasis to enable substitutions at one site to shift the amino-acid preferences of distant sites.

### Entrenchment of substitutions modestly contributes to mutational shifts

One idea that has recently gained support in the protein-evolution field is that substitutions become "entrenched" by subsequent evolution (***Pollock et al., 2012; Shah et al., 2015; Starr et al., 2017***). Entrenchment is the tendency of a mutational reversion to become increasingly unfavorable as a sequence evolves. Given two homologs, if there is no entrenchment then the effect of mutating a site in the first homolog to its identity in the second will simply be the opposite of mutating the site in the second homolog to its identity in the first. But if there is entrenchment, then both mutations will be unfavorable, since the site is entrenched at its preferred identity in each homolog.

Figure 8 shows the distribution of effects for mutating all sites that differ between BG505 and BF520 to the identity in the other Env. As expected under entrenchment, the average effect of these mutations is deleterious - although there are a substantial number of sites where the mutational flips are not deleterious. We can get some sense of the magnitude of the entrenchment by comparing the effects of the BG505↔BF520 mutations to the distribution of effects of all possible amino-acid mutations (Figure 8). This comparison shows that even unfavorable inter-Env mutational flips are generally more favorable than random amino-acid mutations. Therefore, entrenchment occurs for some but not all substitutions that distinguish BG505 and BF520, and the magnitude of entrenchment is less than the effect of a typical random mutation. Entrenchment of substitutions therefore contributes to some of the mutational shifts. But given that many of these shifts occur at sites that do not even differ between the Envs (Figure 7D), entrenchment of substitutions is clearly not the only cause of the shifting amino-acid preferences.

**Figure 8.**
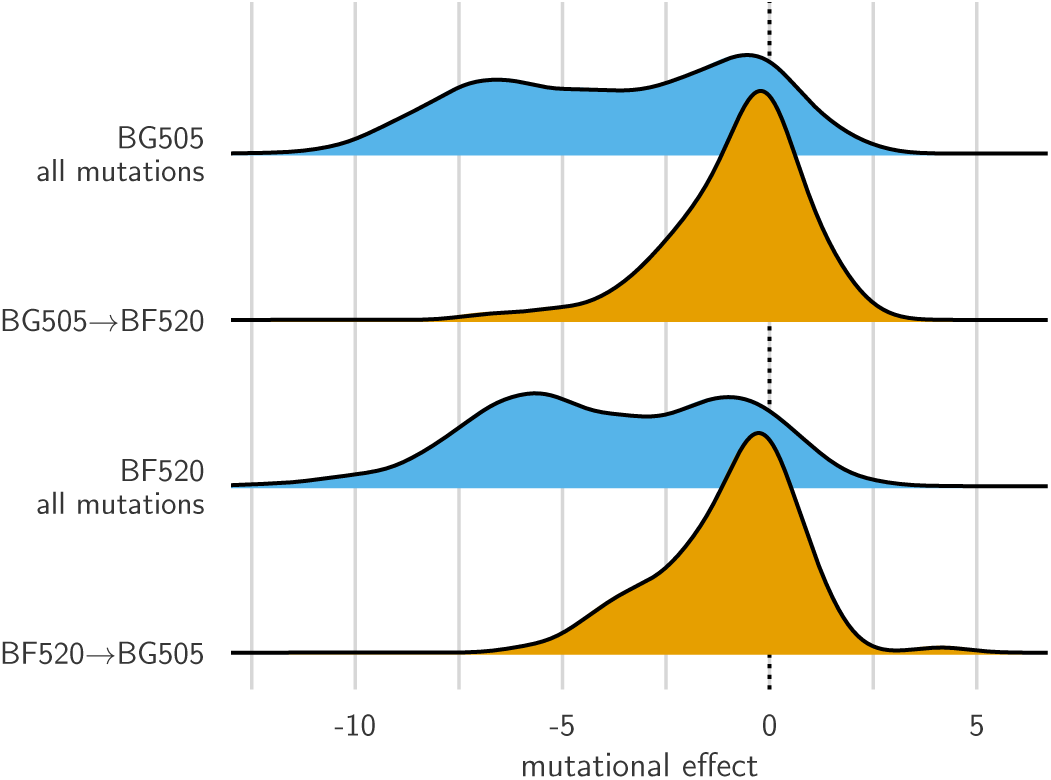
Entrenchment of substitutions during Env evolution. There are 12,521 possible amino-acid mutations at the 659 mutagenized sites alignable between BG505 and BF520. The blue densities show the effects of all these mutations to each Env. The orange densities show the effects of just the 92 mutations that convert BG505 to BF520 or vice versa. In the absence of entrenchment, mutating a site in BG505 to its identity in BF520 should have the opposite effect of mutating the site in BF520 to its identity in BG505. In this case, we would expect the BF520→BG505 distribution to be the mirror image of the BG505→BF520 distribution - and both distributions should be centered around zero if the two Envs are equivalently functional. Instead, mutating a site in either Env to its identity in the other Env tends to be deleterious, indicating that substitutions are often entrenched in the Env in which they have fixed. The effect of a mutation is quantified as the log of the ratio of the site's preference for the mutant amino acid to the preference for the wildtype amino acid.

### Comparing selection in the lab to natural selection

Our experiments measure the effects of mutations on viral growth in a T-cell line in the lab. But HIV actually evolves in humans, where additional selection pressures on Env are undoubtedly present. For instance, antibody pressure might increase the rate of evolution at some sites (***Albert et al., 1990; Wei et al., 2003; Richman et al., 2003***), whereas pressure to mask certain epitopes (***Kwong et al., 2002***) might add constraint at other sites. Comparing selection in our experiments to natural selection can identify sites that are under such additional pressures during HIV's actual evolution in humans.

**Figure 9.**
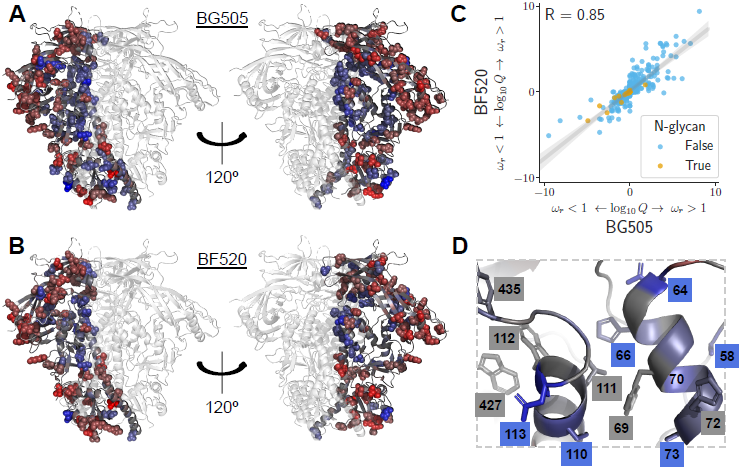
Sites in Env that evolve faster or slower in nature than expected given the functional constraints measured in the lab. We calculated the statistical evidence that each site evolves faster (*ω_r_ >* 1) or slower (*ω_r_* < l)than expected given the experimentally measured amino-acid preferences using the method of ***Bloom (2017)***. **(A)** One monomer of the Env trimer (PDB 5FYL; ***Stewart-Jones et al., 2016***) is colored from blue to gray to red based on the strength of evidence that sites evolve slower than expected (blue), as expected (gray) or faster than expected (red) given the BG505 experiments. Sites where the rate of evolution is significantly different than expected at a false discovery rate of 0.05 are shown in spheres. **(B)** Like (A) but using the data from the BF520 experiments. For both Envs, sites that evolve significantly slower or faster than expected are often on Env's surface (Figure 9-Figure supplement 1). (C) The results are similar regardless of whether the BG505 or BF520 experiments are used. Many of the sites of slower-than-expected evolution are asparagines in N-linked glycosylation motifs (Figure 9-Figure supplement 2). All sites that evolve slower than expected for both experimental datasets are in Figure 9-Figure supplement 3. **(D)** A large cluster of sites that evolve slower than expected is likely involved in Env's transition between open and closed conformational states. Gray boxes indicate sites that ***Ozorowski et al. (2017***, PDB 5VN3) proposed form a hydrophobic network that regulates the conformational change; blue boxes and sticks indicate sites that evolve slower than expected. All analyses used the phylogenetic tree in Figure 1. **Figure 9-Figure supplement 1.** Relative solvent accessibilities of sites evolving faster or slower than expected. **Figure 9-Figure supplement 2.** Amino-acid preferences and alignment frequencies for glycosylation motifs. **Figure 9-Figure supplement 3.** Amino-acid preferences and alignment frequencies of sites that evolve slower than expected. **Figure 9-source data 1.** The *ω_r_* and *Q*-values are in `merged_omegabysite.csv`.

We determined whether each site in Env evolves faster or slower in nature than expected given three models: that evolution is purely neutral (all nonsynonymous and synonymous mutations have equivalent effects), that sites are under the protein-level constraint measured in our experiments with BG505, or that sites are under the constraint measured with BF520. The first model used a standard dN/dS test (the "FEL" method of ***Kosakovsky Pond and Frost, 2005***), whereas the other two models performed a conceptually similar test but used ExpCM's that incorporate experimental measurements as described by ***Bloom (2017)***. The standard dN/dS model finds hundreds of sites that evolve slower than expected under neutral evolution (Table 1, *ω_r_* < 1), and only a handful of sites that evolve faster than expected under neutral evolution (Table 1, *ω_r_* > 1). This finding is unsurprising, since it is well known that Env is under functional constraint. In contrast, ExpCM's that test the rates of evolution relative to the experimentally measured constraints find far fewer sites that evolve slower than expected, but many more sites that evolve faster (Table 1).

The sites that evolve slower or faster than expected from the experiments are shown in Figure 9A,B, and overlaid on the logoplots in Figures 4 and 5 as the *ω_r_* values. The identified sites are similar regardless of whether we use the experiments with BG505 or BF520 (Figure 9C). The reason the results are similar for both experimental datasets is that (as discussed above) the amino-acid preferences of *most* sites are similar in both Envs, suggesting that either dataset provides a reasonable approximation of the site-specific functional constraints across the clade A Envs in Figure 1.

What causes some sites to evolve faster or slower in nature than expected from the experiments? The answer in both cases is likely to be immune selection. Most of the sites of faster-than-expected evolution are on the surface of Env (Figure 9A,B and Figure 9-Figure supplement 1). Env's escape from autologous neutralizing antibodies often involves amino-acid substitutions in surface-exposed regions (***Moore et al., 2009***), including at many of the sites that evolve faster than expected. Since our deep mutational scanning did not impose antibody pressure, sites where substitutions are antibody-driven will evolve faster in nature than expected from the experiments.

Interestingly, immune selection also offers a plausible explanation for the sites that evolve *slower* than expected. In addition to escaping immunity via substitutions at antibody-binding footprints, Env is notorious for employing a range of more general strategies to reduce its susceptibility to antibodies. These strategies include shielding immunogenic regions with glycans (***Wei et al., 2003; Stewart-Jones et al., 2016; Gristick et al., 2016***) or hiding them by adopting a closed protein conformation (***Kwong et al., 2002; Guttman et al., 2015; Ozorowski et al., 2017***). Sites that contribute to such general immune-evasion strategies will be under a constraint in nature that is not present in our experiments - and indeed, such sites evolve more slowly than expected from our experiments. For instance, we find very little selection to maintain most glycans in our cell-culture experiments. Of the 21 N-linked glycosylation sites shared between BG505 and BF520, only four are under strong selection to maintain the glycan in our experiments - despite the fact that most are conserved in nature (Figure 9C and Figure 9-Figure supplement 2). This finding concords with prior literature suggesting that these glycans are selected primarily for their role in immune evasion (***Pugach et al., 2004; Wang et al., 2013; Rathore et al., 2017***). Similarly, a network of sites that help regulate Env's transition between open and closed conformations that have different antibody susceptibilities (Figure 9D) also evolve slower in nature than expected from our experiments. Therefore, we can distinguish evolutionary patterns that are shaped by simple selection for Env function from those that are due to the additional complex pressures imposed during human infections.

## Discussion

We have experimentally measured the preference for each amino acid at each site in the ectodomain and transmembrane domain of two Envs under selection for viral growth in cell culture. These amino-acid preference maps are generally consistent with prior knowledge about sites that are important for protein properties such as receptor binding or disulfide-mediated stability. However, the main value of these maps comes not from comparing them with prior knowledge, but from the fact that such prior knowledge encompasses just a small fraction of the vast mutational space available to Env. Because Env evolves so rapidly, every study of this protein must be placed in an evolutionary context, and our comprehensive amino-acid preference maps potentially enable this in ways that prior piecemeal studies of mutations cannot.

But these maps come with a potentially serious caveat: each one is measured for just a single Env variant. The major question that our study aimed to answer is whether the maps are still useful for evolutionary questions, or whether Env's amino-acid preferences shift so rapidly that each map only applies to the specific HIV strain for which it was measured. This question is reminiscent of one that was grappled with in the early days of protein crystallography, when it first became possible to build maps of a protein's structure. Because it was not (and is still not) possible to crystallize every variant of a protein, it was necessary to determine whether protein structures could be usefully generalized among homologs. Fortunately for the utility of structural biology, it soon became apparent that closely homologous proteins have similar structures (***Chothia and Lesk, 1986; Sander and Schneider, 1991***). This rough generalizability of protein structures holds even for a protein as conformationally complex as Env - for although there are many examples of mutations that alter aspects of Env's conformation and dynamics (***Kwong et al., 2000; White et al., 2010; Almond et al., 2010; Davenport et al., 2013***), SOSIP trimer structures from diverse Env strains remain highly similar in most respects (***Julien et al., 2015; Pugach et al., 2015; Stewart-Jones et al., 2016; Verkerke et al., 2016; Gristick et al., 2016***).

Our results showthatamino-acid preference maps of Env also have a useful level of conservation for many purposes. From a qualitative perspective, the amino-acid preferences look mostly similar between BG505 and BF520, and so provide a valuable reference for estimating which mutations are likely to be tolerated at each site in diverse HIV strains. Indeed, we anticipate that the complete maps of mutational effects in Figures 4 and 5 will be useful for future sequence-structure-function studies. From an analytical perspective, a powerful use of our maps is to identify sites that evolve differently in nature than is required by the simple selection for viral growth imposed in our experiments - and the identified sites are largely the same regardless of whether the analysis uses an amino-acid preference map from BG505 or BF520.

Of course, from the perspective of protein evolution, the most interesting sites are the exceptions to the general conservation of amino-acid preferences. Consistent with studies of other proteins (***Natarajan et al., 2013; Harms and Thornton, 2014; Doud et al., 2015; Starr et al., 2017***), we find a subset of sites that change markedly in which mutations they tolerate. Some shifted sites simply accommodate more amino acids in the more stable BG505 Env - a type of shift that has been well-documented for other proteins (***Wang et al., 2002; Bloom et al., 2006; Gong et al., 2013; Kumar et al., 2017***). But interestingly, there is no strong trend for shifts to be enhanced at sites that differ between BG505 and BF520. Recent studies of protein evolution have focused on the idea that substitutions become "entrenched" as sites shift to accommodate new amino acids (***Pollock et al., 2012; Shah et al., 2015; Bazykin, 2015; Starr et al., 2017***). Indeed, a prior protein-wide comparison of amino-acid preferences across homologs of influenza nucleoprotein found a significant enrichment of shifts at sites of substitutions (***Doud et al., 2015***). But although there is some entrenchment of differences between BG505 and BF520, this is not the major factor behind the shifts in amino-acid preferences: most sites that have shifted between BG505 and BF520 actually have the same wildtype amino acid in both Envs. This rather surprising result might be due to Env's exceptional conformational complexity - mutations can cause long-range alterations in Env's conformation (***Kwong et al., 2000; White et al., 2010; Almond et al., 2010; Davenport et al., 2013***), so it seems plausible that they might also shift mutational tolerance at distant sites.

Our experiments provide highly quantitative data on the mutational tolerance of Env under selection for viral growth in cell culture. These data are amenable to rigorous functional and evolutionary analyses. Here we have shown how these data can be compared between Envs to identify sites where mutational tolerance shifts with viral genotype, or between experiments and nature to identify sites under different pressure in the lab and in humans. Future experiments that modulate selection pressures in other relevant ways should provide further insight into the forces that drive and constrain HIV's evolution.

## Methods and Materials

### Creation of codon-mutant libraries

Our codon mutant libraries mutagenized all sites in *env* to all 64 codons, except that the signal peptide and cytoplasmic tail were not mutagenized. The rationale for excluding these regions is that they are not part of Env's ectodomain, and may be prone to mutations that strongly modulate Env's expression level.

The codon-mutant libraries were generated using the approach originally described in ***Bloom (2014)***, with the modification of ***Dingens et al. (2017)*** to ensure more uniform primer melting temperatures. The computer script used to design the mutagenesis primers (along with some detailed implementation notes) is at https://github.com/jbloomlab/CodonTilingPrimers. For BF520, the three libraries are the same ones described by ***Dingens et al. (2017)***. For BG505, we created three libraries for this study. The wildtype BG505 sequence used for these libraries is in Supplemental file 3. The BG505 mutagenesis primers are in Supplemental file 4.

The end primers for the BG505 mutagenesis were: 5'-tgaaggcaaaactactggtccgtctcgagcagaagacagtggcaatgaga-3' and 5'-gctacaaatgcatataacagcgtctcattctttccctaacctcaggcca-3'. As with BF520, we cloned the BG505 ***env*** libraries into the *env* locus of the full-length proviral genome of HIV strain Q23 (another subtype-A transmitted/founder virus; ***Poss and Overbaugh, 1999***) using the high-efficiency cloning vector described in ***Dingens et al. (2017)***. For this cloning, we digested the cloning vector with BsmBI, and then used PCR to elongate the amplicons to include 30 nucleotides at each end that were identical in sequence to the ends of the BsmBI-digested vector. The primers for this PCR were: 5'-agataggttaattgagagaataagagaaagagcagaagacagtggcaatgagagtgatgg-3' and 5'-ctcctggtgctgct ggaggggcacgtctcattctttccctaacctcaggccatcc-3'. Next, we used NEBuilder HiFi DNA Assembly (NEB, E2621S) to clone the *env* amplicons into the BsmBI-digested plasmids. We purified the assembled products using Agencourt AMPure XP beads (Beckman Coulter, A63880) using a bead-to-sample ratio of 1.5, and then transformed the purified products into Stellar electrocompetent cells (Takara, 636765). The transformations yielded between 1.5-3.6 million unique clones for each of the three replicate libraries, as estimated by plating 1:2,000 dilutions of the transformations. We scraped the plated colonies and maxiprepped the plasmid DNA; unlike in ***Dingens et al. (2017)***, we did not include a 4-hour outgrowth step after the scraping step. For the wildtype controls, we maxiprepped three independent cultures of wildtype BG505 *env* cloned into the same Q23 proviral plasmid. See Figure 2-Figure supplement 1 and Figure 3A for information on the average mutation rate in these libraries as estimated by Sanger sequencing and deep sequencing, respectively.

### Generation and passaging of viruses

For BG505, we generated mutant virus libraries from the proviral plasmid libraries by transfecting 293T cells in three 6-well plates (so 18 wells total per library) with a per-well mixture of 2 μg plasmid DNA, 6 *μ*l FuGENE 6 Transfection Reagent (Promega, E269A), and 100 *μ*l DMEM. The 293T cells were seeded at 5 × 10^5^ cells/well in D10 media (DMEM supplemented with 10% FBS, 1% 200 mM L-glutamine, and 1% of a solution of 10,000 units/mL penicillin and 10,000 *μ*g/mL streptomycin) the day before transfection, such that they were approximately 50% confluent at the time of transfection. In parallel, we generated wildtype viruses by transfecting one 6-well plate of 293T cells with each wildtype plasmid replicate. At 2 days post-transfection, we harvested the transfection supernatant, passed it through a 0.2 *μ*m filter to remove cells, treated the supernatant with DNAse to digest residual plasmid DNA as in ***Haddox et al. (2016)***, and froze aliquots at −80°C. We thawed and titered aliquots using the TZM-bl assay in the presence of 10 *μ*g/mL DEAE-dextran as described in ***Dingens et al. (2017)***.

We conducted the low MOI viral passage illustrated in Figure 2A in SupT1.CCR5 cells (obtained from Dr. James Hoxie; ***Boyd et al., 2015***). During this passage, cells were maintained in R10 media, which has the same composition as the D10 described above, except RPMI-1640 (GE Healthcare Life Sciences, SH30255.01) is used in the place of DMEM. In addition, the media contained 10 *μ*g/mL DEAE-dextran to enhance viral infection. We infected cells with 4 million (for replicate 1) or 5 million (for replicates 2 and 3) TZM-bl infectious units of mutant virus at an MOI of 0.01, with cells at a starting concentration of 1 million cells/mL in vented tissue-culture flasks (Fisher Scientific, 14-826-80). At day 1 post-infection, we pelleted cells, aspirated the supernatant, and resuspended cell pellets in the same volume of fresh media still including the DEAE-dextran. At 2 days post-infection, we doubled the volume of each culture with fresh media still including DEAE-dextran. At 4 days post-infection, we pelleted cells, passed the viral supernatant through a 0.2 *μ*m filter, concentrated the virus ∼30 fold using ultracentrifugation as described in ***Dingens et al. (2017)***, and froze aliquots at −80°C. In parallel, for each replicate, we also passaged 2 × 10^5^ (for replicate 1) or 5 × 10^5^ (for replicates 2 and 3) TZM-bl infectious units of wildtype virus using the same procedure. To obtain final titers for our concentrated virus, we thawed one of the aliquots stored at −80°C and titered using the TZM-bl assay in the presence of 10 ywg/mL DEAE-dextran.

For the final short-duration infection illustrated in Figure 2A, for each replicate we infected 10^6^ TZM-bl infectious units into 10^6^ SupT1.CCR5 cells in the presence of 100 *μ*g/mL DEAE-dextran (note that this is a 10-fold higher concentration of DEAE-dextran than for the other steps, meaning that the effective MOI of infection is higher if DEAE-dextran has the expected effect of enhancing viral infection). Three hours post-infection, we pelleted the cells and resuspended them in fresh media without any DEAE-dextran. At 12 hours post-infection, we pelleted cells, washed them once with PBS, and then used a miniprep kit to harvest reverse-transcribed unintegrated viral DNA (***Haddox et al., 2016***).

The generation, passaging and deep sequencing of BF520 was done in a highly similar fashion, except that we only had a single replicate of the wildtype control. Note that the final passaged BF520 mutant libraries analyzed here actually correspond to the "no-antibody" controls described in ***Dingens et al. (2017)***, but that study did not analyze the initial plasmid mutant libraries relative to these passaged viruses, and so was not able to provide measurements of the amino-acid preferences.

### Illumina deep sequencing

We deep sequenced all of the samples shown in Figure 3A: the plasmid mutant libraries and wildtype plasmid controls, and the cDNA from the final mutant viruses and wildtype virus controls. In order to increase the sequence accuracy, we used a barcoded-subamplicon sequencing strategy. This general strategy was originally applied in the context of deep mutational scanning by ***Wu et al. (2014)***, and the specific protocol used in our work is described in ***Doud and Bloom (2016)*** (see also https://jbloomlab.github.io/dms_tools2/bcsubamp.html).

The primers used for BG505 are in Supplementary file 5. The primers used for BF520 are in ***Dingens et al. (2017)***. The data generated by the Illumina deep sequencing are on the Sequence Read Archive under the accession numbers provided at the beginning of the Jupyter notebook in Supplementary files 1 and 2.

### Analysis of deep-sequencing data

We analyzed the deep-sequencing data using the `dms_tools2` software package (***Bloom, 2015***, https://jbloomlab.github.io/dms_tools2/). The algorithm that goes from the deep-sequencing counts to the amino-acid preferences is that described in ***Bloom (2015)*** (see also https://jbloomlab.github.io/dms_tools2/prefs.html). A Jupyter notebook that performs the entire analysis including generation of most of the figures in this paper is in Supplementary file 1. An HTML rendering of this notebook is in Supplementary file 2.

The Jupyter notebooks in Supplementary files 1 and 2 also contain numerous plots that summarize relevant aspects of the deep sequencing such as read depth, per-codon mutation frequency, mutation types, etc. Supplementary file 1 also contains text files and CSV files with the numerical values shown in these plots.

Citations are also owed to `weblogo` (***Crooks et al., 2004***, http://weblogo.threeplusone.com/) and `ggseqlogo` (***Wagih, 2017***, https://omarwagih.github.io/ggseqlogo/), which were used in the generation of the logoplots.

### Alignments and phylogenetic analyses of Env sequences

A basic description of the process used to generate the clade A sequence alignment in Figure 1 - source data 1, the alignment mask in Figure 1 -source data 2, and the phylogenetic tree in Figure 1 are provided in the legend to that figure. An algorithmic description of how the alignment and tree were generated are in Supplementary files 1 and 2.

For fitting of the phylogenetic substitution models, we used phydms (***Hilton et al., 2017***, http://jbloomlab.github.io/phydms/) to optimize the substitution model parameters and branch lengths on the fixed tree topology in Figure 1. The Goldman-Yang (or YNGKP) model used in Table 1 is the M5 variant described by ***Yang et al. (2000)***, with the equilibrium codon frequencies determined empirically using the CF3×4 method (***Pond et al., 2010***). For the ExpCM shown in Table 1, we extended the models with empirical nucleotide frequencies described in ***Hilton et al. (2017)*** to also allow *ω* to be drawn from discrete gamma-distributed categories exactly as for the M5 model. These ExpCM with gamma-distributed *ω* were implemented in phydms using the equations provided by ***Yang (1994)*** (see also http://jbloomlab.github.io/phydms/implementation.html#models-with-a-gamma-distributed-model-parameter). The preferences were re-scaled by the stringency parameters in Table 1 as described in ***Hilton et al. (2017)***. For both the M5 model and the ExpCM with a gamma-distributed *ω*, we used four categories for the discretized gamma distribution.

Table 1 -source-data 1 shows the results for a wider set of models than those used in Table 1. These include the M0 model of ***Yang et al. (2000)***, ExpCM without a gamma-distributed *ω*, and ExpCM in which the amino-acid preferences are averaged across sites as a control to ensure that the improved performance of these models is due to their site-specificity. Note how for these Env alignments, using a gamma-distributed *ω* is very important in order for the ExpCMs to outperform the M5 model - we suspect this is because there are many sites of strong diversifying selection.

For detection of sites with faster or slower than expected evolution, we used the approach in ***Bloom (2017)***, which is exactly modeled on the FEL approach of ***Kosakovsky Pond and Frost (2005)*** but extended to ExpCM. This approach estimates a ***P***-value that *ω_r_* is not equal to one for each site *r* using a likelihood-ratio test. For the Q-values and false discovery rate testing, we considered the tests for *ω_r_* > 1 and *ω_r_* < 1 separately.

Supplementary files 1 and 2 contains the code that runs phydms to reproduce all of these analyses.

### Identifying sites of shifted amino-acid preference

When identifying shifts in amino-acid preferences between the two Envs, we needed a way to quantify differences between the Envs while accounting for the fact that our measurements are noisy. The approach we use is based closely on that of ***Doud et al. (2015)***, and is illustrated graphically in Figure 6A. The RMSD_corrected_ value is our measure of the magnitude of the shift. Figure 6A, its legend, and the associated text completely explains these calculations with the following exception: they do not detail how the "distance" between any two preference measurements was calculated. The distance between preferences at each site was simply defined as half of the sum of absolute value of the difference between preferences for each amino acid. Specifically, for a given site *r*, let 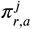 be the preference for amino-acid *a* in homolog *i* (e.g., BG505) and let 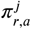 be the preference for that same amino acid in homolog *j* (e.g., BF520). Then the distance between the homologs at this site is simply 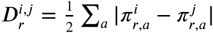. The factor of 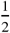 is used so that the maximum distance will always fall between zero and one.

### Analysis of entrenchment

For the analysis in Figure 8, the results are presented in terms of the mutational effects rather than the amino-acid preferences. If *π_r,a_* is the preference of site *r* for amino-acid *a* and *π_ra_* is the preference for amino-acid *a′* (both re-scaled by the stringency parameters in Table 1), then the estimated effect of the mutation from *a* to *a′* is simply log 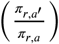.

### Data and code availability

All code and input data required to reproduce all analyses in this paper are in Supplementary file 1 (see also Supplementary file 2). The deep sequencing data are on the Sequence Read Archive with the accession numbers listed in Supplementary files 1 and 2.

## Acknowledgments

We thank Michael Doud and Orr Ashenberg for computer code that formed the basis for some of the analyses. We thank Andrew Ward for pointing out to us that some of the sites with slower-than-expected rates of evolution are involved in Env's conformational changes upon receptor binding. We thank Kelly Lee for helpful input about the relative stabilities of the BG505 and BF520 Envs. We thank the Fred Hutch Genomics Core for performing the Illumina deep sequencing. This work was supported by the following grants from the NIH: DP1-DA039543 from the NIDA to JO and R01-AI127893 from the NIAID to JDB. The work was also supported in part by a Collaboration for AIDS Vaccine Discovery Grant (OPP1111923). The research of JDB is supported in part by a Faculty Scholar Grant from the Howard Hughes Medical Institute and the Simons Foundation. HKH was supported by a Cell and Molecular Biology Training Grant (T32GM007270) from the NIGMS of the NIH. ASD was supported by an NSF Graduate Research Fellowship (DGE-1256082).

---

**Supplementary file 1.** The code to perform all steps in the analysis is in `analysis_code.zip`. Specifically, this file contains a Jupyter notebook that performs the analysis, all required input data, and all reasonably sized output files. The Jupyter notebook downloads the deep sequencing data, processes it with the `dms_tools2` software (***Bloom, 2015***, https://jbloomlab.github.io/dms_tools2/), and also performs a variety of downstream analyses that generate most of the figures for this paper.

**Supplementary file 2.** An HTML rendering of the Jupyter notebook that performs the computational analysis. The actual notebook is in Supplementary file 1, but if you just want to look at the analysis rather than run it, then you may prefer this file instead. In particular, the notebook contains plots detailing the deep sequencing data analysis as generated using the `dms_tools2` software (***Bloom, 2015***, https://jbloomlab.github.io/dms_tools2/).

**Supplementary file 3.** The sequence of the wildtype BG505 *env* used in our study is in FASTA format in the file `BG505_env.fasta`.

**Supplementary file 4.** The sequences of the primers used for the BG505 codon mutagenesis are in the file `BG505_codon_mutagenesis_primers.txt`.

**Supplementary file 5.** The primers used for the BG505 barcoded-subamplicon sequencing are in the file `BG505_bcsubamp_primers.txt`.

**Figure 1-Figure supplement 1.**
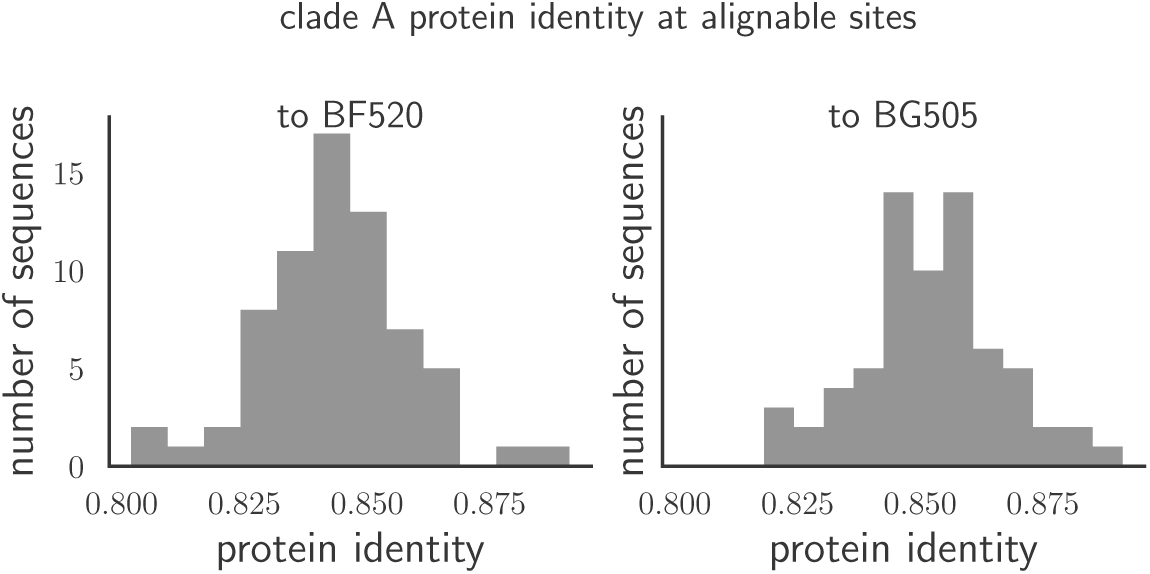
The histograms show the pairwise amino-acid identity of each Env to all other sequences in the clade A alignment in Figure 1 -source data 1 after masking the sites delineated in Figure 1-source data 2. There are 616 non-masked sites. The pairwise protein identity between BG505 and BF520 is 86.2% (721 of 836 sites identical) when considering ***all*** sites, and 89.1% (549 of 616 sites identical) when considering just the non-masked sites.

**Figure 2-Figure supplement 1.**
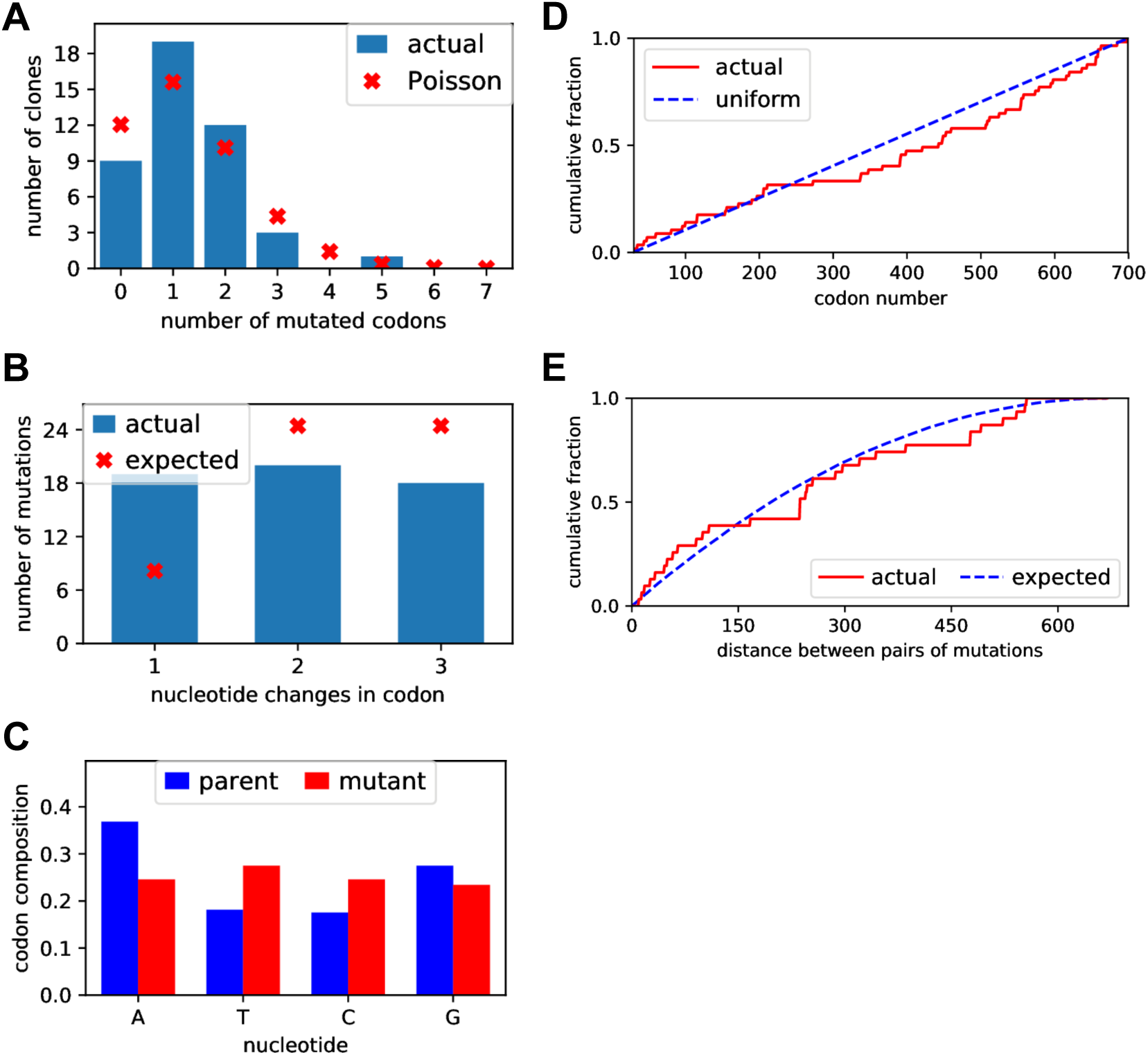
We Sanger sequenced 44 clones of BG505 Env sampled roughly evenly from each of the three replicate mutant plasmid libraries. **(A)** There was an average of 1.5 mutant codons per clone, with the number of mutations per clone roughly following a Poisson distribution. **(B)** The mutant codons had a mix of single-, double-, and triple-nucleotide changes, with an elevated number of single-nucleotide changes than expected. **(C)** Nucleotide frequencies were fairly uniform in the mutant codons. **(D)** Mutations were distributed roughly evenly along the mutagenized region of *env* (30-699 in the sequential numbering scheme used in this plot). **(E)** For clones with multiple mutations, we computed the pairwise distance in primary sequence between each codon mutation and plotted the cumulative distribution of these distances (red line). We also simulated the expected distribution of pairwise distances if mutations occurred entirely independently (blue line). The observed distribution is close to the expected distribution. Comparable data for the BF520 libraries is provided in ***Dingens et al. (2017)***.

**Figure 3-Figure supplement 1.**
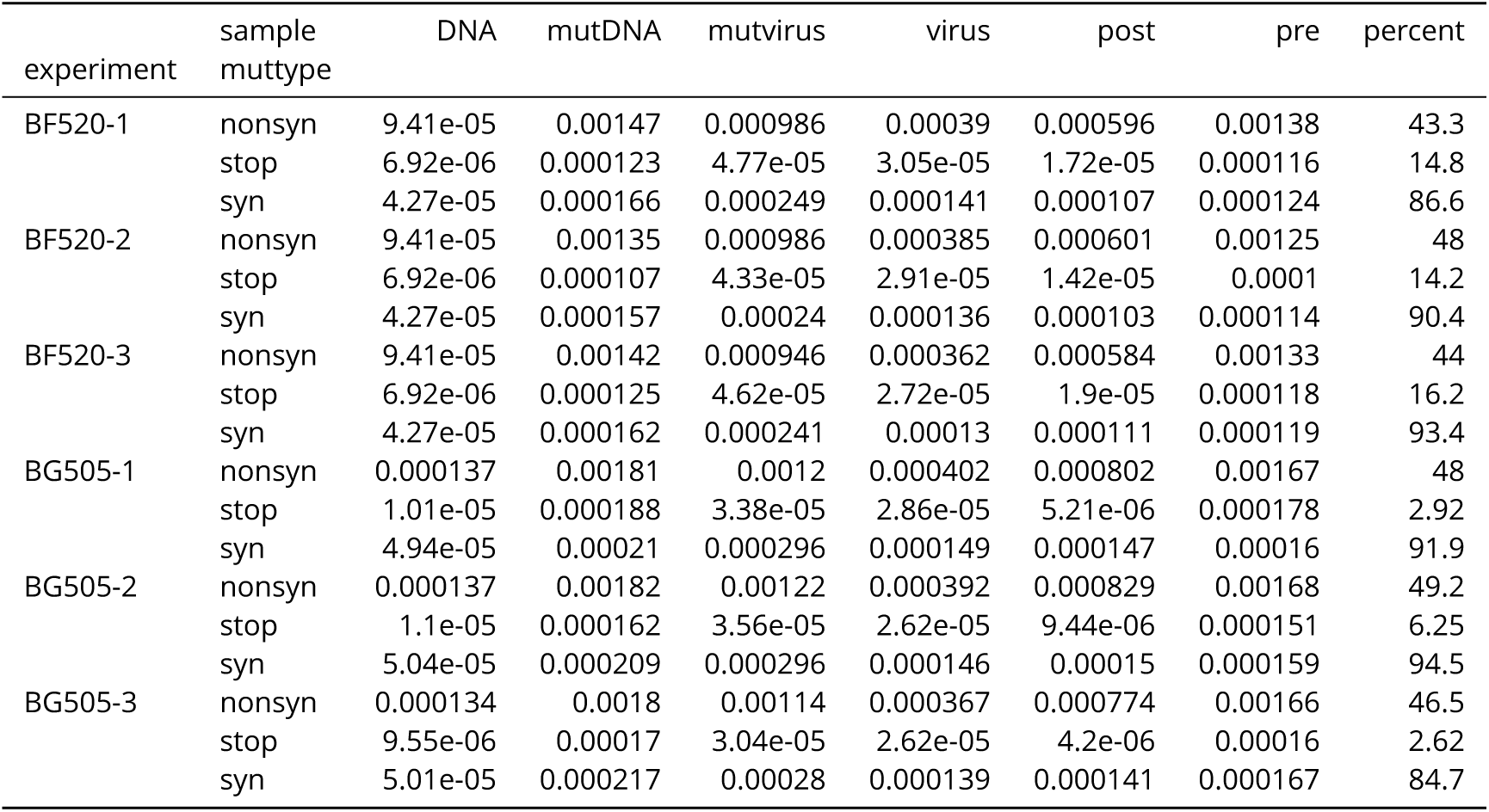
This table gives the average frequencies of nonsynonymous, synonymous, and stop-codon mutations as plotted in Figure 3. Note that there is only one *DNA* sample for BF520, so that same sample is repeated three times in the sample in conjunction with each BF520 replicate. We calculate the error-corrected pre-selection mutation frequency as the *mutDNA* frequency minus the *DNA* frequency, and the error-corrected post-selection mutation frequency as the *mutvirus* frequency minus the *virus* frequency. We then use these error-corrected frequencies to calculate the *percent* of mutations remaining after selection for each replicate and type of mutation.

**Figure 7-Figure supplement 1.**
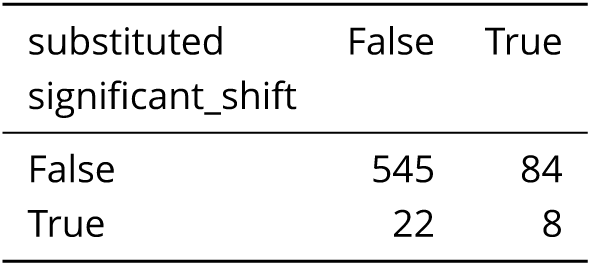
The sites of significant shifts as determined in Figure 6B are somewhat more likely to be among the sites that have substituted between BG505 and BF520. However, this association is only borderline statistically significant, with ***P*** = 0.055 using a Fisher's exact test.

**Figure 9-Figure supplement 1.**
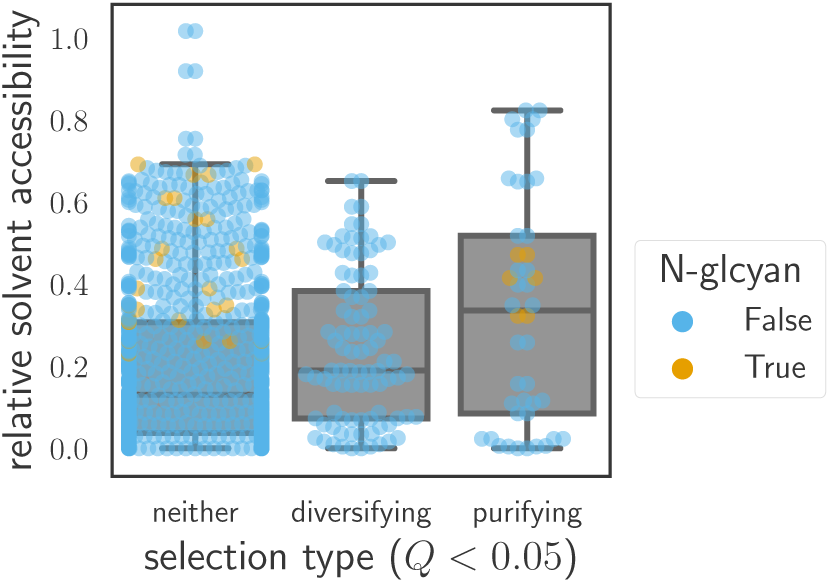
Sites are grouped by whether they have ***œ_r_ > 1*** (diversifying selection) or *ω_r_* < 1 (purifying selection) at ***Q*** < 0.05 in *both* Envs, or whether they fall into neither of these categories. Relative solvent accessibilities were calculated as in Figure 7. As can be seen from these box plots with overlaid points for each site, sites of both diversifying and purifying selection tend to have higher relative solvent accessibility than other sites.

**Figure 9-Figure supplement 2.**
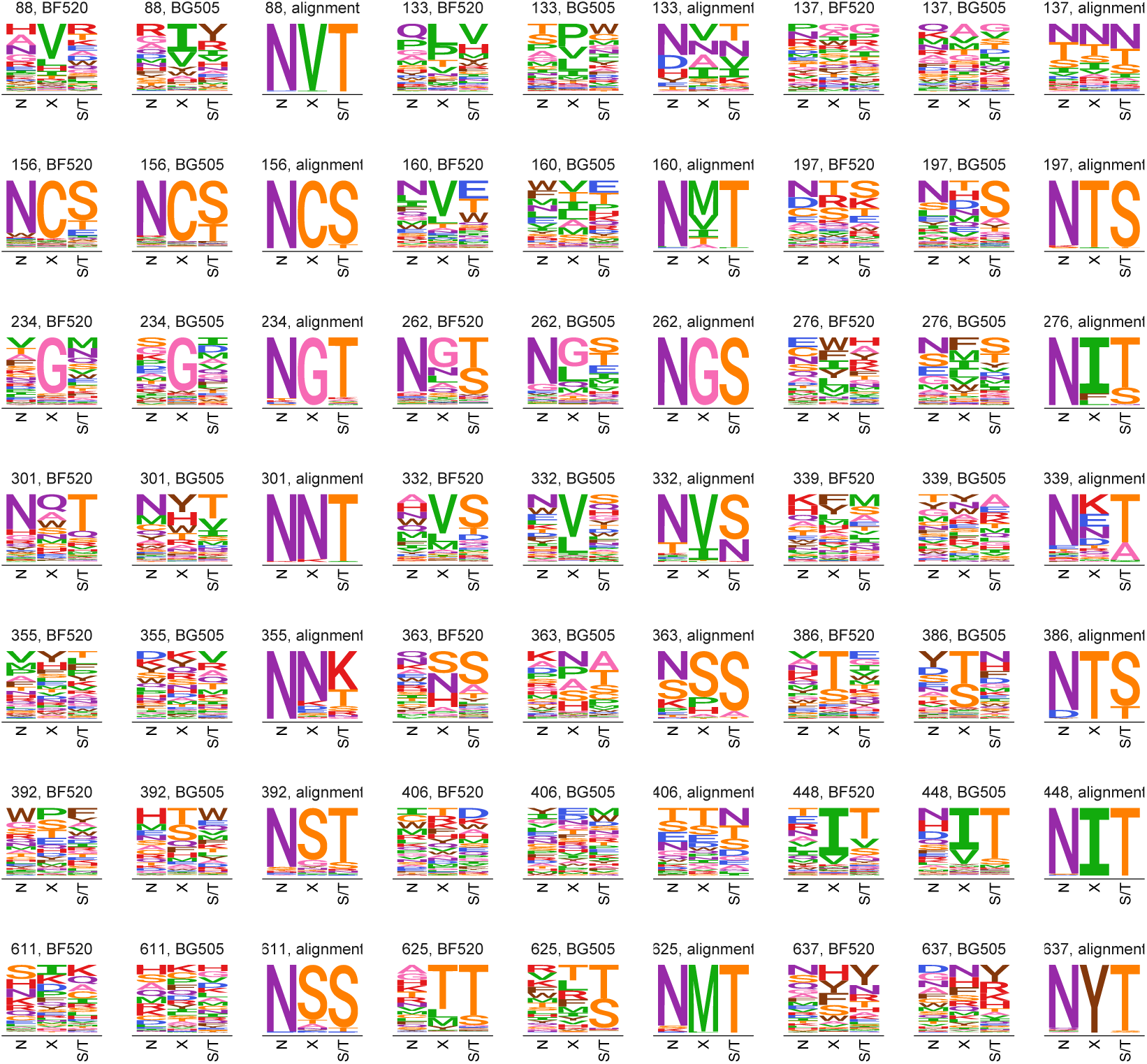
This plot shows all sites that are N-linked glycosylation motifs in both BG505 and BF520. Each motif is named by the residue number of the asparagine, and the preferences (averaged across replicates) for each Env are shown as well as the frequencies of amino acids across the clade A alignment at each of the three positions in the motif (N-X-S/T). As can be seen from this plot, the experiments measure relatively broad mutational tolerance at many sites where the natural Env sequences have a strongly conserved motif. We suspect this is because many glycans serve as a shield against immunity in nature - a function that is not required in cell culture.

**Figure 9-Figure supplement 3.**
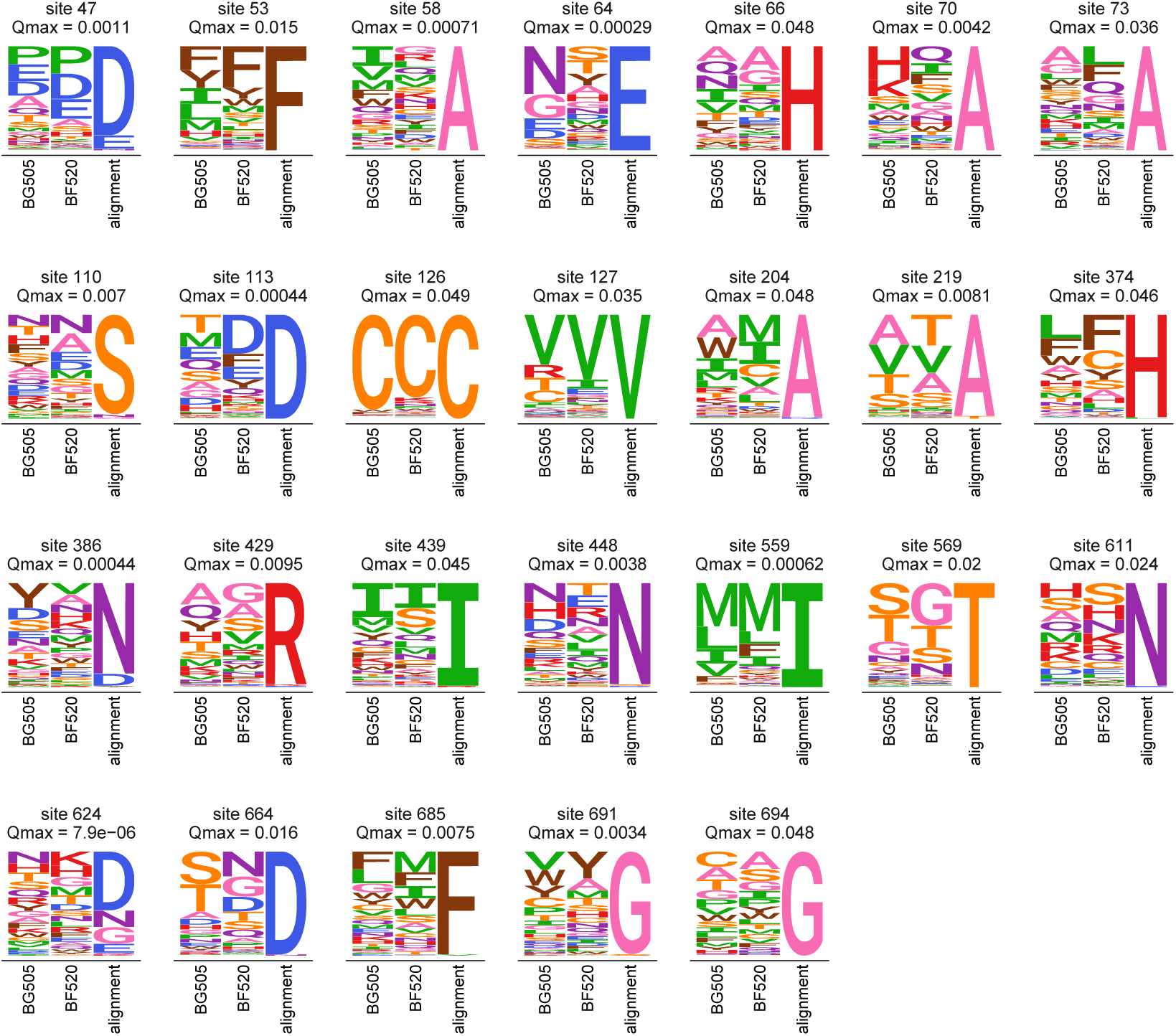
This plot shows all sites that are evolving more slowly than expected in natural sequences given the preferences measured in both Envs. Specifically, it shows all sites with *ω_r_* < 1 at ***Q*** < 0.05 for the ExpCMs for *both* BG505 and BF520. For each site, the plots show the preferences averaged across replicates and re-scaled for each Env, as well as the frequencies of amino acids in the clade A Env alignment. The Q-value indicated is the maximum of that for the two Envs.

